# Effective expression analysis using gene interaction matrices and convolutional neural networks

**DOI:** 10.1101/2021.09.07.459284

**Authors:** Arvind Pillai, Piotr Grabowski, Bino John

## Abstract

Artificial intelligence recently experienced a renaissance with the advancement of convolutional neural networks (CNNs). CNNs require spatially meaningful matrices (*e.g*., image data) with recurring patterns, limiting its applicability to high-throughput omics data. We present GIM, a simple, CNN-ready framework for omics data to detect both individual and network-level entities of biological importance. Using gene expression data, we show that GIM-CNNs can outperform comparable neural networks in performance and their design facilitates network-level interpretability. GIM-CNNs provide a means to discover novel disease-relevant factors beyond individual genes and their expression, factors that are likely missed by standard differential gene expression approaches.

## Introduction

Convolutional neural network (CNN) architectures have emerged as one of the most powerful approaches in machine learning^1,2^. The use of CNN-centric architectures enables automatic inference of relevant features from image-based data and allows data scientists to leverage the rich array of methodologies in both CNNs and, more broadly, deep learning (DL). However, due to the vectorized nature of omics data, generic frameworks to analyze expression data using CNNs has been limited with the added caveat of loss of interpretability with the best performing methods. For example, DeepInsight (DI) converts gene-expression data to an image-like feature matrix using unsupervised dimensionality reduction (t-SNE, k-PCA) and image processing^3^. DI uses complex data transformations using latent features to yield the relevant image matrices for CNNs, making it a powerful tool. However, such complex transformations also make it nearly impossible to provide a biologically meaningful interpretation of the resulting dimensionality-reduced features. A probabilistically-grounded framework to predict gene relationships from single cell sequencing data using a co-expression based approach^4^ that derives a normalized empirical probability distribution function (NEPDF) for CNNs was also recently developed. NEPDFs are powerful as additional inputs to CNNs to predict gene-gene relations. NEPDFs currently rely on processing of very large sets of relevant data such as single cell sequencing to build the underlying NEPDF matrices. More recently, several different CNN architectures were tested^5^ to predict cancer types using RNA-Seq data from TCGA^6^. The resulting observations further substantiates previous studies on the advantages of using CNNs for gene expression. However, the end-users are restricted by the CNN-architectures and are limited to classifying cancer types as implemented in the code. Due to the restricted availability and limitations of methods available for applying CNNs on gene-expression data, we developed a novel, generalized approach, termed GIM (Gene Interaction Matrices). GIM uses a biologically inspired, gene-interaction based data transformation on gene expression data to create an image-like feature matrix from any gene expression-based study. The transformed data can then be used with any CNN-based machine learning approach for a variety of challenging problems such as disease diagnostics and drug development. We compared the performances of a number of standard neural network architectures including GIM-CNNs on two disparate datasets^6,7^ to illustrate the utility of GIM-CNNs in classification problems. We further use kidney cancer as an example to illustrate the ability of GIMs to unravel disease relevant interactions. We show that GIM-CNNs can identify important differentially regulated genes, as well as complex gene-gene links that are non-trivial to uncover using standard differential gene expression (DEG) techniques.

## Methods

Detailed information on datasets, feature selection, and testing strategies are described in the supplementary material. GIM is designed to make use of high-throughput readouts of defined entities such as those of genes (*e.g*., gene expression data) from the primary samples of interest (*e.g*., treatment samples), and from the samples that represent the corresponding baseline biological signals, such as “control” samples. For example, given two study groups of a gene expression experiment such as treated vs control samples, we denote the corresponding gene expression data by 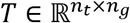, and 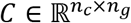, where *n_t_*, *n_c_*, and *n_g_* are number of treatment replicates, number of control replicates, and genes of interest (*e.g.*, most variant genes). GIM uses the gene expression datasets to compute a square matrix (*A*) as the input for a CNN, comprising of a harmonic gene-gene score (diagonal and lower triangular matrix), and a relative gene-gene expression score (upper triangular matrix). Specifically, given two genes *g_i_*, and *g_j_*, the cell value, *A_ij_* for a sample is calculated as shown (equations 1-3).

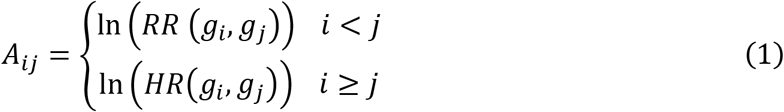

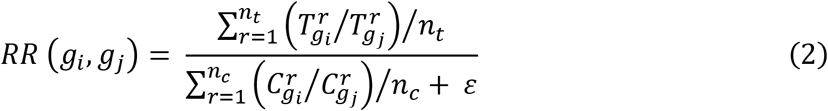

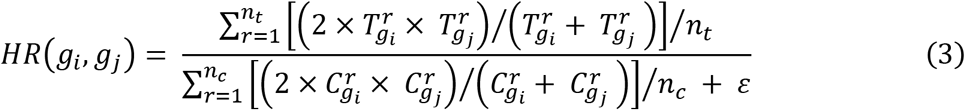

Where *i, j* = 1 … *n_g_*, *ε* = 1 × 10^-5^, 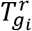 and 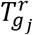 represents value of genes *g_i_* and *g_j_* in the *r*-th treatment replicate, respectively; 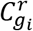 and 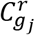 represents value of genes *g_i_* and *g_j_* in the *r*-th control replicate, respectively. Note that for diagonal elements of *A* (*i.e*. *i*=*j*), the harmonic mean ratio (*HR*) scores reduces to the equivalent of fold-change values for each gene, allowing CNNs to explicitly use this important feature. *HR* scores thus capture the changes in average cumulative abundance of gene pairs between two experimental conditions. In contrast, relative ratio (RR) scores measure the changes in the relative abundance of gene pairs (*e.g*., perturbations in interdependent pathways). Replacement of genes by other biological entities of interest in the equations would allow for the application of GIMs to other similar data such as proteomics and metabolomics. GIM is made available for free through the AstaZeneca R&D Github (https://github.com/AstraZeneca/GIM).

## Results

We tested GIM in two different scenarios (Figure 1a): (1) a binary classification problem with matched treatment and control samples using the Open TG-Gates (TGG) data^7^, and (2) a more intricate, multi-class classification problem on TCGA datasets with the added complexity of not having an experimentally defined, matched controls^6^. GIM was first applied to the TGG dataset of 160 compounds using a CNN (GIM-CNN) to predict if a given compound-induced gene expression perturbation, is adverse. GIM-CNN yielded the highest MCC^8^ scores of 0.42 over all different neural network-based architectures tested, corresponding to an improvement of ~10-20% in performance over the other methods. Even with the use of thousands more features (7400 vs 225 genes) the next best performing method, CNN-Re^5^ only yielded an MCC of 0.38. For the second test using the TCGA cancer dataset, we used GIM-CNN to predict 33 cancer types. Once again, GIM-CNN performed better than all other methods tested when using an identical feature set of 225 genes. GIM also enabled highly comparable classification accuracy to that of CNN-Re (93.10 vs 93.8 %), while using only a small proportion (3%) of the features used by CNN-Re^5^. Finally, since each GIM feature represents a pseudo measure of biologically relevant interactions, GIM-CNNs presents a direct route to discover potentially important regulators, interactions, and networks (Fig. 1C). For example, the top 30 important gene interactions for Kidney Renal Clear Cell Carcinoma naturally, and perhaps surprisingly, fall into a well-connected network with UPK1B, GPX2, and AQP4 representing the top three hubs, all of which have been linked to renal cancer. For instance, the top connected gene, UPK1B has recently emerged as an essential gene for renal urothelium function^9^, and is a known promoter of bladder metastasis^10^. GPX2, a renal cancer-promoting gene, manifests high expression in specific clinical sub-types only^11^, highlighting the feature of GIM-CNNs in extracting meaningful links that are challenging for typical DEG-based approaches. Finally, AQP4, belonging to the aquaporin family, has an established role in cancer biology^12^ with clinical links to bladder cancer^13,14^, suggestive of an underappreciated role for AQP4 in renal cancer biology.

**Figure 1:**
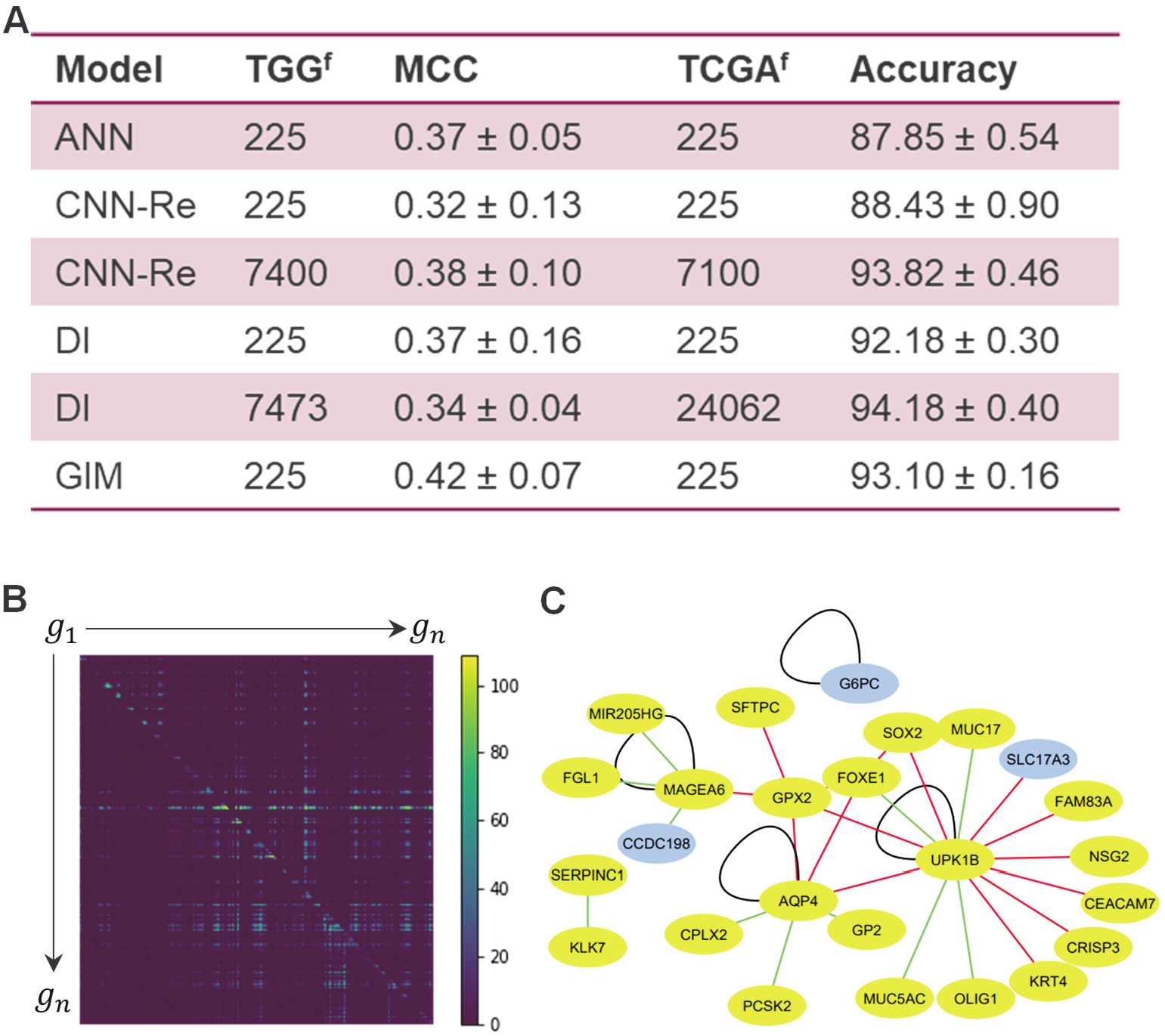
**A)** Performance comparisons of various methods on TGG and TCGA datasets (See supplement for details). ANN: Artificial neural network, CNN-Re: CNN-reshape, DI: DeepInsight, MCC: Matthew’s correlation coefficient; Accuracy: Categorical Accuracy. TGG^f^ and TCGA^f^ represent number of features used for the TGG, and TCGA datasets, respectively. **B)** Visualization of the GIM feature importance map for Kidney Renal Clear Cell Carcinoma (KIRC) using GRAD-CAM. Brighter color corresponds to most important GIM features (gene pairs) for KIRC classification. **C)** Top 30 most important features (gene pairs) extracted from the feature importance map (Fig 1b), spanning 25 unique genes. Edge colors correspond to types of feature interactions. Green: harmonic mean score (HR, lower triangle of the matrix), black: gene log2 fold-change (HR scores at diagonal of the matrix), red: relative ratio score (RR, upper triangle of the matrix). Known KIRC biomarkers are marked as blue nodes.

In summary, GIM-CNNs presents several other advantages over other machine learning methods for gene expression data analysis, simultaneously enabling network-level biological discoveries and strong predictive performance while using modest number of features. In contrast to non-DL methods, GIM allows for leveraging the rich and complex information hidden in a high-dimensional gene expression dataset. With respect to other DL methods such as DI for gene expression, GIM also provides the advantage of having a deterministic feature encoding step. These attributes of GIM should improve reproducibility and interpretability in machine learning workflows using CNNs. The proposed framework can be easily also applied to similar high-dimensional vectorized data including but not limited to epigenome profiling (*e.g*., ATAC-Seq, ChIP-Seq), and proteome profiling (*e.g*., mass-spec proteomics), and metabolomics. Since CNN’s can make use of highdimensional tensors, GIM framework could also enable additional data about gene-gene relationships (*e.g*., PPI, proteomics data) in future studies. GIM may also prove useful in leveraging the powerful transfer learning approach (*e.g.,* ResNet^15^) to address machine learning problems where large training datasets are not available. For instance, GIM-CNNs could be pretrained on large datasets such as TCGA and single cell sequencing, and subsequently fine-tuned with smaller datasets of interest for predictive as well as hypothesis generation needs.

## Acknowledgements

Sincere thanks to AstraZeneca (AZ) internal reviewers, Nigel Greene (AZ), and Mishal Patel (AZ).

## Conflict of Interest

none declared.

## Notes

### Competing Interest Statement

The authors have declared no competing interest.

