## Supplementary Materials for "Effective expression analysis using gene interaction matrices and convolutional neural networks"

### Supplementary information

### 1. Datasets

#### 1.1. TG-Gates (TGG) rat *in-vivo* toxicogenomics data

Microarray-based Open TG-Gates rat liver *in-vivo* dataset[1] (single- and repeat-dosing experiments) was retrieved from the hosting institution's FTP server (ftp://ftp.biosciencedbc.jp/archive/open-tggates/). The CEL files were RMA-normalized using the R 'oligo' package version 1.52.1 with 'normalize' and 'background' arguments set to TRUE. Probesets mapping to the same gene were aggregated using the 'max' function. The mapping table between Rat Genome 230 2.0 probesets and Ensembl gene IDs was obtained from Affymetrix website (https://www.affymetrix.com/, file Rat230_2.na36.annot.csv.zip). A recent study[2] using the TGG data was used to establish the target labels (Disease States, DS) for method comparisons. These DS are linked to adverse liver events in rats given specific compounds in various doses and time-points, while non-DS examples are linked to drug treatments which had no toxic effects in rat livers. Disease States not directly relevant to liver toxicity (DS9, kidney injury) or not having a unique gene expression signature (DS3, hemolysis and DS4, low TBIL) were removed. All Disease States were pooled into one positive class while non-DS treatments were used as the negative class. Consequently, 833 positive and 2,683 negative examples were used in this study. Log2 array intensity was used as the gene expression unit in the machine learning tasks.

#### 1.2. The Cancer Genome Atlas (TCGA) transcriptomics data

The Cancer Genome Atlas (TCGA) RNAseq data[3] was downloaded using the R TCGAbiolinks package version 2.16.4[4]. The data from 33 different cancer types was downloaded in the raw HTSeq count format. Genes with mean raw counts <10 were filtered out and the Variance-Stabilising Transformation (VST) from DESeq2 version 1.28.1[5] was applied to the filtered dataset. The 33 cancer types were modelled as a multi-class classification task. Additionally, the normal tissue samples in TCGA were removed resulting in 10,363 examples and 24,062 features (genes). TCGA data sets did not have replicates and matched controls were unavailable for many samples. Therefore, a pseudo-control sample was used by averaging over all samples within each set (training, validation, and test sets).

### 2. Feature Processing

#### 2.1. Feature Selection

Prior to any method-specific dimensionality reduction, the microarray TGG dataset was reduced to 50% of its original feature space by removing genes correlated with each other by more than 0.5 Pearson correlation coefficient, reducing the number of features to 7473. Furthermore, the top 225 variable genes (based on their coefficient of variance, $c_v = \sigma / \mu$) were selected for the ANN, CNN-Re, DI and GIM experiments with both TGG and TCGA datasets. The features were limited to 225 to minimize memory needs for CNNs and the same number of features was then used for all algorithms to enable fair assessment of performance. In the TGG dataset, the $c_v$ was calculated across all dose timepoints within each compound treatment group and the genes most constantly varying across the different compound groups were used as the top 225 for the entire TGG dataset. For TCGA samples, the entire dataset was used for $c_v$ calculation.

#### 2.2. Additional feature formatting for the CNN-Re and DeepInsight benchmarks

To apply CNN-Re (CNN-Reshape[6]), the 1D gene expression vector of each sample was reshaped into a rectangular image-form matrix. The DeepInsight (DI) method[7] reduces the features based on a dimensionality reduction technique to form an image representation of the gene expression vectors. The DeepInsight (https://github.com/alok-ai-lab/DeepInsight) open-source package was used to transform the 1D gene expression vectors into 64 × 64 image-like feature matrices using default DI t-SNE (2 components and cosine distance as the metric, perplexity set to 30.0). In addition to the direct 225-feature comparison of CNN-Re and DI methods to our GIM approach, CNN-Re and DI runs using larger feature sets were also tested. The number of features used for CNN-Re was 7,400 in case of the TGG data and 7,100 in case of the TCGA dataset. This was done to replicate the original CNN-Re approach which used 7100 features in the TCGA cancer prediction problem (out of 24,062 total features). For the TGG dataset, which was not originally used in the CNN-Re publication, a similar number of features was kept to compare the change of performance of CNN-Re when changing from 225 to a high-dimensional feature space. Similarly, the additional DI experiments with larger feature sets were performed to replicate the original DI approach which allows to transform a very high-dimensional feature set using t-SNE or k-PCA to an image-like matrix. For the DI experiment on the TGG data, the entire filtered TGG dataset was used (7473 features) while for the TCGA dataset, all 24062 features were employed to create the 64x64 image matrices for convolutional neural networks.

### 3. Training and cross-validation

#### 3.1 TGG dataset

The dataset was split into five folds and trained using 32-compounds-out cross-validation to minimize training data leak, common in biological machine learning tasks. Matthew's Correlation Coefficient (MCC) was used to measure the performance, and an early-stopping criteria parameter 'patience' was set to 5 on the validation MCC. As the goal here was to predict if a compound-dose-timepoint is toxic or not, the binary cross-entropy loss function and the Adam optimizer were used. All the models were optimized based on architectures and parameters as described section 4. The best training hyperparameters and architectures for each Model presented in Figure 1. B (main paper) are described below (Table 1)

| Model | Architecture | Learning rate | Batch size | Epochs |
|---|---|---|---|---|
| ANN | ANN | $1 \times 10^{-4}$ | 64 | 150 |
| CNN-Re | Shallow CNN | $1 \times 10^{-5}$ | 64 | 100 |
| DI | ResNet | $1 \times 10^{-5}$ | 64 | 100 |
| GIM | Shallow CNN | $1 \times 10^{-5}$ | 64 | 40 |

**Table 1. Best training hyperparameters and model-architecture pairs in the TGG dataset**

#### 3.2 TCGA dataset

The TCGA dataset was split into training and hold-out sets with 8,290 and 2,073 samples, respectively. Next, the training was evaluated using 5-fold stratified cross-validation to predict cancer subtype as a multi-class classification problem. This ensured the same proportion of labels in the training and validation folds. The models trained on each fold using the categorical cross-entropy loss function, the Adam optimizer, and an early-stopping criteria with patience = 7 was set on the validation accuracy within the fold. Subsequently, the trained folds were tested on the hold-out and their results are reported in Figure 1B (main manuscript). The best training hyperparameters and architectures for each model are described below (Table 2).

| Model | Architecture | Learning rate | Batch size | Epochs |
|---|---|---|---|---|
| ANN | ANN | $1 \times 10^{-4}$ | 64 | 150 |
| CNN-Re | Shallow CNN | $1 \times 10^{-5}$ | 128 | 200 |
| DI | ResNet | $1 \times 10^{-5}$ | 128 | 200 |
| GIM | Shallow CNN | $1 \times 10^{-5}$ | 128 | 200 |

**Table 2. Best training hyperparameters and model-architecture pairs in the TCGA dataset**

### 4. Neural network architectures

#### 4.1. Artificial Neural Network (ANN).

The ANN implemented contained four fully connected (FC) layers as shown in Table 3. Every FC layer, except the output layer, is batch-normalized and activated with a ReLU function. The number of neurons and dropout percentage considered for tuning were [512, 256, 128, 64, 32, 16] and [0.3, 0.4, 0.5], respectively. The best combination is presented in Table 3 below. Additionally, the selected values for tuning the learning rates, batch sizes, and epochs were [$1 \times 10^{-3}, 1 \times 10^{-4}, 1 \times 10^{-5}$], [64, 128, 256], and [50, 80, 100, 120, 150], respectively.

| Layer | Output Shape |
|---|---|
| FC1 | 128 |
| FC2 | 64 |
| Dropout1 (0.5) | 64 |
| FC3 | 32 |
| Dropout2 (0.5) | 32 |
| FC4 | 16 |
| FC5 (Output) | 1 or 33 |

**Table 3. Final ANN architecture.**

#### 4.2. Shallow CNN.

A shallow CNN architecture (**Table 4**) using the Vanilla-CNN architecture[6] was used for CNN-Re and the proposed method. The numbers of filters, kernel sizes, learning rate, batch size, and epochs selected for tuning were [16, 32, 64], [3, 5, 10], [$1 \times 10^{-3}, 1 \times 10^{-4}, 1 \times 10^{-5}$], [64, 128, 256], and [40, 80, 100, 150, 200], respectively. Moreover, we performed a network architecture search by tuning the convolution block size. A convolution block is a single or set of 2D convolutional layers followed by a maxpooling layer.

| Layer | Output Shape | Kernel | Filters (strides) |
| --- | --- | --- | --- |
| Convolution 2D | $216 \times 216$ | $10 \times 10$ | $16\ (1)$ |
| Maxpooling 2D | $108 \times 108$ | $2 \times 2$ | N/A (1) |
| Flatten | 373248 | N/A | N/A |
| FC1 | 128 | N/A | N/A |
| FC2 (output) | 1 or 33 | N/A | N/A |

**Table 4. Shallow CNN - final architecture used for CNN-Re and GIM-CNN approaches.**

#### 4.3. ResNet

A Residual Network with 4 layers of 3 residual blocks each from the basic residual in [8] was implemented (**Table 5**) and used with the 64x64 images created by DeepInsight. A custom grid search was used to optimize the ResNet architecture by tuning the number of residual layers and residual blocks with [3, 4, 5] and [2, 3, 4, 5], respectively. Additionally, the learning rate, batch size, and epochs chosen for tuning were [$1 \times 10^{-4}, 1 \times 10^{-5}$], [64, 128, 256], [50, 100, 150, 200].

| Layer | Output size | Kernel size | Filters (stride) |
| --- | --- | --- | --- |
| Convolution 2D | $64 \times 64$ | $7 \times 7$ | $16\ (1)$ |
| Maxpooling 2D | $32 \times 32$ | $2 \times 2$ | $N/A\ (2)$ |
| residual convolution1 | $32 \times 32$ | $\begin{Bmatrix} 3 \times 3 \\ 3 \times 3 \end{Bmatrix} \times 3$ | $\begin{Bmatrix} 16\ (1) \\ 16\ (1) \end{Bmatrix} \times 3$ |
| residual convolution2 | $16 \times 16$ | $3 \times 3$ $\begin{Bmatrix} 3 \times 3 \\ 3 \times 3 \end{Bmatrix} \times 3$ | $32\ (2)$ $\begin{Bmatrix} 32\ (1) \\ 32\ (1) \end{Bmatrix} \times 3$ |
| residual convolution3 | $8 \times 8$ | $3 \times 3$ $\begin{Bmatrix} 3 \times 3 \\ 3 \times 3 \end{Bmatrix} \times 3$ | $64\ (2)$ $\begin{Bmatrix} 64\ (1) \\ 64\ (1) \end{Bmatrix} \times 3$ |
| residual convolution4 | $4 \times 4$ | $3 \times 3$ $\begin{Bmatrix} 3 \times 3 \\ 3 \times 3 \end{Bmatrix} \times 3$ | $128\ (2)$ $\begin{Bmatrix} 128\ (1) \\ 128\ (1) \end{Bmatrix} \times 3$ |
| Global average pooling | 128 | $N/A$ | $N/A$ |
| FC | $1\ or\ 33$ | $N/A$ | $N/A$ |

**Table 5. ResNet architecture used with DeepInsight transformation.**

### 5. Performance metrics

#### Matthews Correlation Coefficient (MCC):

MCC is defined as a function of True Positives (TP), True Negatives (TN), False Positives (FP), and False Negatives (FN).

$$MCC = \frac{TP \times TN - FP \times FN}{\sqrt{(TP + FP)(TP + FN)(TN + FP)(TN + FN)}}$$

#### Accuracy:

Categorical accuracy was used which calculates the percentage of predicted values that match with the actual values for the one-hot vector (33 cancer subtypes in TCGA). Assuming both true and predicted label vectors are one-hot encoded, a correct prediction is recorded if the index of the maximal true value is equal to the index of the maximal predicted value.

$$Accuracy = \frac{Number\ of\ correct\ predictions}{Total\ number\ of\ samples} \times 100$$

### 6. Interpretability

GIM-CNN enabled us to directly gain important biological insights from the gene expression data using CNN interpretability techniques. We used GRAD-CAM[9] using the tf-keras-vis package (https://github.com/keisen/tf-keras-vis) to the image-like feature matrices of a cancer subtype in the test set to generate importance values. For every Gene Interaction Matrix, the continuous importance values were converted to binary maps by considering only the 99$^{th}$ percentile values in the harmonic gene-gene score (diagonal and lower triangular elements) and the relative gene-gene expression score (upper triangular elements), separately to minimize biases due to any variance differences from the two different scores. Important gene pairs for a cancer subtype were subsequently identified by summing these binary maps to generate a heatmap. Finally, the most important gene-pairs were extracted from the heatmap to investigate if they form a well-connected network. For example, the heatmap for Kidney Renal Clear Cell Carcinoma (KIRC) over 108 test samples is shown Figure 1B in the main manuscript; and Figure 1C shows that the top 30 most important gene-pairs form a well-connected network of biological relevance, using Cytoscape[10].
